# Proteomic Response of *Thalassiosira pseudonana* to Anoxia Reveals Alanine Fermentation Pathway and Reprogramming of Nitrogen Metabolism

**DOI:** 10.64898/2026.07.20.739533

**Authors:** Gwenaëlle Gain, Malika Chabi, Nicolas Berne, Hervé Degand, Javier Cordoba, Pierre Morsomme, Fabrice Bray, Claire Remacle, Ugo Cenci, Pierre Cardol

## Abstract

- Diatoms are key players in marine ecosystems and are frequently exposed to low-oxygen conditions in sediments and oxygen minimum zones. However, the metabolic strategies that enable their survival under anoxia remain poorly understood.
- Using the model diatom *Thalassiosira pseudonana,* we investigated its response to dark anoxia. We first used proteomic approaches to identify differences between dark anoxia and dark oxic conditions combined with a phylogenetic analysis to decipher how the anoxic tolerance was acquired by this lineage. This approach revealed unexpected shunts involving amino acid pathways. We then correlated these findings with targeted metabolomic analysis on amino acids.
- Our results show that the diatom *T. pseudonana* undergoes a coordinated metabolic reprogramming under anoxia, centered on alanine production and tightly coupled to nitrogen metabolism. This work reveals how carbon and nitrogen fluxes are integrated to maintain cellular homeostasis in the absence of oxygen and provides a framework for understanding the resilience of diatoms in oxygen-depleted environments.
- We identify three key features for anoxic adaptation in this lineage (i) alanine-centered metabolic reprogramming as a central component of acclimation to anoxia (ii) the involvement of an arginine-succinate shunt; and (iii) a critical contribution of lateral gene transfer (LGT) for anoxic tolerance.

## Introduction

Hypoxic and anoxic environments are widespread in marine ecosystems, particularly in coastal sediments and expanding oxygen minimum zones. Microalgae that persist in these environments must maintain metabolic activity despite the absence of oxygen, yet the underlying mechanisms remain poorly understood (Banti *et al*., 2013; Catalanotti *et al*., 2013). Diatoms are among the most abundant and ecologically important phytoplankton groups, playing a central role as primary producers and major contributors to global oxygen production (Field *et al*., 1998; Bar-On *et al*., 2018; Falciatore *et al*., 2020; José *et al*., 2026). Diatoms, including members of genera *Thalassiosira* and *Cyclotella,* are also abundant in turbid estuarine waters and even in oxygen-depleted sediments (Muylaert *et al*., 2006; Broman *et al*., 2017).

Despite their prevalence in these environments, physiological studies on diatoms under hypoxic or anoxic conditions remain scarce. As a result, the molecular mechanisms that enable diatoms to survive and maintain metabolism during oxygen deprivation are still largely unknown.

Several species, including *Thalassiosira weissflogii*, have been shown to store nitrate intracellularly and use it as an alternative electron acceptor (Ramírez *et al*., 1966; Lomas *et al*., 2000) through dissimilatory nitrate reduction to ammonium (DNRA) providing a means to sustain redox balance under oxygen limitation (Kamp *et al*., 2015, 2016; Stief *et al*., 2022; Stenow *et al*., 2024). In addition, genomic analyses have revealed that diatoms encode a diverse set of enzymes potentially involved in fermentation metabolism (Atteia *et al*., 2013). Notably, the genome of the centric diatom *Thalassiosira pseudonana* encodes a remarkably diverse set of enzymes, including a [FeFe]-hydrogenase, a pyruvate:ferredoxin oxidoreductase (PFO), a pyruvate formate lyase (PFL), an acetate succinate-CoA transferase (ASCT), and an ADP-dependent acetyl-CoA synthetase (ADP-ACS). However, the functional relevance and integration of these pathways under anoxia remain largely unresolved.

In other photosynthetic eukaryotes, such as the green alga *Chlamydomonas reinhardtii*, anaerobic metabolism has been extensively characterized and involves multiple fermentative pathways that sustain glycolytic flux from starch degradation (Mus *et al*., 2007). In brown algae and other photosynthetic Stramenopiles, a functionally similar role could be fulfilled by laminarin, the main storage polysaccharide, whose mobilization provides hexoses that feed glycolysis and, more broadly, central metabolism (Graiff *et al*., 2016; Chen *et al*., 2021), particularly under low-light conditions and possibly in response to certain environmental stresses (Roessler, 1988; Murison *et al*., 2024). This would maintain redox balance in the absence of active mitochondrial respiratory chain and tricarboxylic acid (TCA) cycle (Mus *et al*., 2007; Matthew *et al*., 2009; Catalanotti *et al*., 2012, 2013; Atteia *et al*., 2013). In contrast, the metabolic strategies that support energy production and redox homeostasis in diatoms are still unclear. Addressing this question requires a detailed characterization of metabolic and regulatory responses. Although the model diatom *T. pseudonana* has been widely used in molecular and ecological studies, proteomic investigations have so far focused mainly on other environmental stressors, such as pollutant exposure(Gain *et al*., 2023), nutrient limitation (Du *et al*., 2014), or CO₂ fluctuations (Clement *et al*., 2017).

Here, we investigated the acclimation of *T. pseudonana* to dark anoxia by combining quantitative proteomics with targeted amino acid quantification. We showed that anoxia induces a coordinated reorganization of central metabolism, in which alanine production emerges as a major metabolic node linking carbon and nitrogen metabolism. A phylogenomic analysis further revealed the evolutionary context of these responses within diatoms. Together, our results uncover a previously unrecognized anoxic response in *T. pseudonana*, involving glutamine- and aspartate-based metabolic routes, and suggest that part of this capacity may derive from lateral gene transfer (LGT).

## Materials and Methods

### Strain and culture conditions

The axenic strain of Thalassiosira pseudonana (CCMP 1335), kindly provided by A. Falciatore and B. Bailleul (IBPC, Paris, France), was grown under low photosynthetic photon flux density (PPFD, low light, 50 μmol photons.m−2.s−1), using white light-emitting diode (LED), under 12 h light/12 h dark cycle at 18 °C. Cultures were maintained in artificial seawater at a salinity of 33 g.L−1, supplemented with f/2 medium and silica (Sigma-Aldrich, G9903) as described in Guillard and Ryther (1962) (Guillard & Ryther, 1962) and Guillard (1975) (Guillard, 1975). Experiments were performed with cells in exponential growth. Cell concentrations were measured using a Coulter Z2 Particle Counter (Beckman, Indianapolis, IN, USA) with ∼4 µm size threshold. Methods to achieve anoxic conditions. Liquid cultures were centrifuged for 4 min at 4,000 x g, and pellets were resuspended in fresh f/2 medium to a final concentration of 106 cells.mL−1. Cells were acclimated in the dark for 30 min before induction of anoxia. All steps were carried out at room temperature (22 ± 2°C). Cell suspensions were transferred into Erlenmeyer flasks placed into a custom-built sealed chamber (<0.1 μM O2), where cultures were bubbled with nitrogen gas (N2) in the dark. Control (oxic) cultures were maintained under identical conditions but bubbled with air (21% O2).

### Proteomic experiment

#### Sampling and protein extraction

Cell suspensions (106 cells mL−1, 350 mL) were incubated under anoxia and sampled at 0, 1, 2, 4, and 24 h. All subsequent steps were performed at 4°C. Cells were centrifuged (4 min at 4,000 x g) and resuspended in 1 mL extraction buffer (280 mM Mannitol, 10 mM MOPS-KOH, 0.5 mM PMSF). Glass beads (500 µL, 1 mm diameter; Sigma-Aldrich) were added, and cells were lysed using a TissueLyser II (Qiagen) at 30 Hz for 10 10 min (two cycles). Cellular debris were removed by sequential centrifugation at 2,000 x g (10 min) and 5,000 x g (4 min). Protein concentration was quantified by Bradford proteins assay (Biorad) using a Bovine Serum Albumin (BSA) standard curve. Protein precipitation, solubilization and digestion. Proteins were precipitated using the methanol/chloroform method (Wessel & Flügge, 1984) as described in Gain et al., 2021 (Gain et al., 2021). Dry protein pellets were solubilized in 20 µL of 50 mM triethylammonium bicarbonate (TEAB, pH 8.0) containing 0.5 % Rapigest surfactant (Waters) via sonication (Bioruptor, Diagenode). Proteins were reduced with 25 mM tris (2-carboxyethyl) phosphine at 60°C during 1 h and alkylated with 200 mM methyl methanethiosulfonate at RT for 15 min in the dark. The samples were diluted fivefold with a 50 mM TEAB buffer to achieve a final RapiGest concentration of 0.1 %. Proteins were digested for 16 h at 37°C with sequencing-grade modified trypsin (Promega) at a protease-to-protein ratio of 1:20. RapiGest was degraded by incubating samples with 0.5 % trifluoroacetic acid (TFA) f at 37 °C for 1h. After centrifugation (54,000 rpm, 4°C, TLAA55, Optima-Beckman), the supernatant was collected, further centrifuged for 20 min, and dried under vacuum (SC 200 Savant Speed Vac concentrator).

Nano-UPLC Analysis. Dried peptides were resuspended 20 µL of 0.1 % (v/v) formic acid with 2 % (v/v) acetonitrile (ACN). Peptides were separated on a NanoACQUITY UPLC MClass system (Waters ®) controlled by MassLynx V4.1 software (Waters®). Peptides were first trapped on a C18 trap column (100Å, 5 µm, 180 µm x 20 mm, Waters®), desalted using isocratic conditions with at a flow rate of 8 µL min−1 using a 95 % formic acid and 5 % (v/v) ACN buffer for 3 min, and then separated on a C18 analytical column (100 Å, 1.8 µm, 75 µm x 150 mm column, PepMap Waters®) over 130 min at 35°C at flow rate of 300 nL min−1 using a two-part linear gradient from 1 % (v/v) ACN, 0.1% formic acid to 35 % (v/v) ACN, 0.1 % formic acid and from 35 % (v/v) ACN, 0.1 % formic acid to 85 % (v/v) ACN, 0.1 % formic acid.The nano-UPLC was coupled online to the mass spectrometer via a nano-electrospray ionization (nanoESI) source emitter.

#### Ion Mobility Separation-High Definition Enhanced (IMS-HDMSE) analyses

IMS-HDMSE analyses were performed using a SYNAPT G2-Si mass spectrometer (Waters) equipped with a NanoLockSpray dual electrospray ion source (Waters). Precut fused silica PicoTipR Emitters (360 µm OD; 20 µm ID; 10 µm tip; 2.5” length; Waters) were used for sample spraying and identical emitters without tips were used for lock-mass injection. The spray voltage was 2.4 kV, with a sampling cone voltage of 25 V and source offset of 30 V; the source temperature was 80°C. Data were acquired in positive and resolution mode from 15 min to 106 min post injection, in the m/z range 50-2000, using a 1s scan time. Collision energy was ramped from ion mobility bin 20 (20 eV) to 110 (45 eV): low-. The collision energy in the transfer cell for low-energy MS mode was set to 4 eV. Lock-mass correction was performed using [Glu1]-fibrinopeptide B (100 fmol. µL−1) infused every 30 s at 0.5 µL.min−1.

### Data processing for label free quantification

HDMSE data were processed with Progenesis QI software (Nonlinear DYNAMICS, Waters) using T. pseudonana ASM14940v2 reference proteome. Carbamidomethylation (Cys) was set as a fixed modification and Met oxidation as a variable modification; trypsin digestion with one miss cleavage allowed. Four biological replicates were analyzed per condition. Relative quantification used the non-conflicting peptide method. Protein abundances were averaged across replicates, and Log2 Fold Changes (LFCs) values computed for Oxic vs Control (OvsC), Anoxic vs Control (AvsC), and Anoxic vs Oxic (AvsO). Proteins were considered differentially accumulated with ≥3 unique peptides and ANOVA p < 0.05. Heatmaps and hierarchical clustering were generated using the heatmap R package (v1.0.12) with correlation-based distance metrics and centroid linkage. K-means clustering was also applied to identify temporal response patterns.

### Phylogenetic analysis and protein localization

Protein sequences identified in the proteomic dataset were used as queries in BLAST searches against the NCBI non-redundant (nr) database and additional resources (MMETSP (Keeling et al., 2014), JGI). Up to 1000 homologs with E-value < 1e-10 were retrieved and aligned using MAFFT (Katoh & Standley, 2013) with the quick alignments settings. Block selection was performed using BMGE (Criscuolo & Gribaldo, 2010) with a block size of 4 and the BLOSUM30 matrix. Preliminary phylogenetic trees were generated using Fasttree (Price et al., 2010), and sequence redundancy was reduced by ‘dereplication’ of well-supported monophyletic groups using TreeTrimmer (Maruyama et al., 2013). Final alignments were generated with MUSCLE (Edgar, 2004), and block selection was carried out using BMGE with the same settings as above. Phylogenetic trees were inferred with IQ-TREE using the LG model with 1000 ultrafast bootstraps. Trees were visualized using Figtree (64). In addition, The prediction of the 65 proteins identified to upregulated where submitted to targetP 2.0 prediction (https://services.healthtech.dtu.dk/services/TargetP-2.0/) (Emanuelsson et al., 2007; Armenteros et al., 2019) and ASAFind 2.0 (a prediction tools dedicated to indicate localization of proteins from diatoms and other organisms derived from secondary plastid endosymbiosis) (https://asafind.jcu.cz) (Gruber et al., 2015; Gruber & Kroth, 2024; Gruber & Oborník, 2024) using recommended setting.

### Amino Acid Analysis by Dansyl Chloride Derivatization and LC–MS/MS

The composition of the 20 major amino acids was analyzed using a dansyl chloride–specific derivatization method followed by liquid chromatography–tandem mass spectrometry (LC–MS/MS). Cell pellets corresponding to 10⁶ cells were resuspended in 500 µL of methanol 100% and transferred to tubes containing 10 mg of 0.7-mm beads. Samples were disrupted by bead beating with two cycles of 8 min agitation separated by a 20 min incubation period. Following centrifugation, supernatants were collected and dried using a SpeedVac concentrator at 30 °C. Dried extracts were resuspended in 50 µL of 100 mM ammonium bicarbonate. An aliquot of 25 µL was mixed with 12.5 µL of acetonitrile (ACN) and 12.5 µL of 100 mM ammonium bicarbonate buffer (pH 8.8) and vortexed thoroughly. Dansyl derivatization was initiated by adding 25 µL dansyl chloride solution (8 mg/mL in ACN), followed by vortex mixing. Samples were incubated for 1 h at 40 °C. The reaction was quenched by adding 5 µL of 250 mM NaOH to neutralize excess dansyl chloride, followed by vortex mixing and an additional incubation for 10 min at 40 °C. Subsequently, 25 µL of 425 mM formic acid prepared in a 50:50 (v/v) ACN/water solution was added, and the samples were vortexed and evaporated to dryness using a SpeedVac. Dried samples were reconstituted in 100 µL of water and filtered using a 96-well plate equipped with a 0.45 µm PVDF membrane to remove aggregates. Prior to filtration, wells were preconditioned with 100 µL of 70% ethanol and washed twice with 200 µL of Milli-Q water. Filtrates were collected without further drying.

LC–MS/MS analysis was performed using a standard 1-h proteomics method. The filtrate was diluted 1:50 in water containing 0.1% formic acid to avoid signal saturation, and 1 µL of the diluted sample was injected into the LC–MS/MS system.

LC–MS/MS analysis was performed on nanoHPLC U3000 RSLC nanoESI Q-Exactive plus (ThermoScientific). The filtrate was diluted 1:50 in water containing 0.1% formic acid to avoid signal saturation, and 1 µL of the diluted sample was injected into the LC–MS/MS system. One microlitre of dansyl amino acid was injected with solvent A (5% acetonitrile and 0.1% formic acid v/v) for 3 min at a flow rate of 5 μL·min−1 on an Acclaim PepMap100 C18 pre-column (5 μm, 300 μm i.d. × 5 mm) from ThermoFisher Scientific.

The amino acids were then separated on a C18 Acclaim PepMap100 C18 reversed phase column (3 μm, 75 μm i.d. × 500 mm) using a linear gradient (5%–40%) of solution B (75% acetonitrile and 0.1% formic acid) at a rate of 250 nL·min−1. The column was washed with 100% of solution B during 5 min and then re-equilibrated with buffer A. The column and the pre-column were placed in an oven at a temperature of 45°C. The entire analysis was completed in approximately 140 min. The LC runs were performed in positive ion mode with MS scans from m/z 310 to 1400 in the Orbitrap mass analyser. PRM method was achieved with MS resolution 70k, AGC 1e6, max IT 80 ms with mass range m/z 310-1400. For MS/MS, resolution was 17,5K, AGC 5e5, max IT 200 max. The selection windows was m/z 1.8 and HCD with nce 30. Dansyl amino acid m/z was selected :301.1406, 304.2477, 327.0077, 329.0046, 355.0693, 356.0692, 357.0658, 371.315, 372.3183, 374.0971, 391.2835, 445.1198, 446.1201, 447.1163, 447.3464, 448.1165, 462.146, 463.1461, 519.1385, 520.1391. LC–MS/MS data were processed by Quant Browser (Thermo).

Data were analyzed with RStudio using one-way ANOVA, with p-values adjusted for multiple testing by the Benjamini–Hochberg FDR procedure; thresholds (p < 0.05, p < 0.001) refer to FDR-adjusted p-values

## Results

### Global proteomic overview reveals a delayed but extensive response to anoxia

To characterize the response of *Thalassiosira pseudonana* to oxygen deprivation, we performed a time-resolved quantitative proteomics analysis under dark anoxic or oxic conditions for up to 24h, using light-oxic conditions as reference. Across all conditions, 928 proteins were identified (Supplementary Information Table S1). Among these proteins, 58% were annotated, 33% were uncharacterized (*i.e.* of unknown function), and 9% were predicted from genomic data without experimental validation. Annotated proteins spanned a broad range of major cellular functions, including amino acid metabolism, photosynthesis, transcription/translation, and oxidative stress response (Fig. 1a), indicating that anoxia induces global metabolic remodeling.

**Figure 1.**
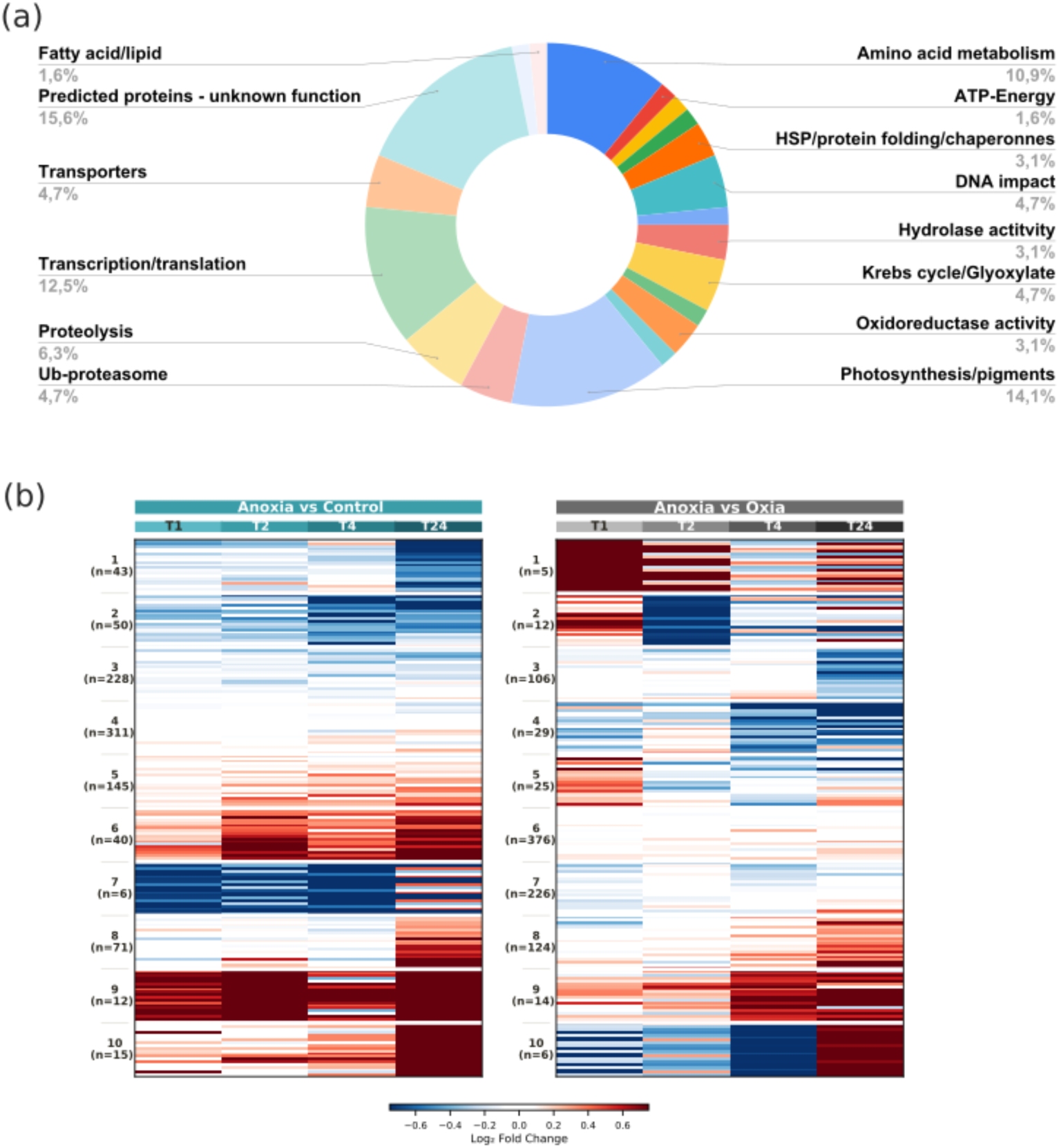
Overview of the proteomic dataset for *Thalassiosira pseudonana* under anoxia. **(a)** Distribution of log_2_ fold change (LFC) for the 928 identified proteins after 24h of anoxia relative to the light-oxic control (A*vs*C, 24h). The histogram shows a symmetric distribution centered near zero, with proteins displaying both increased and decreased abundance. **(b) Clustering analysis of protein abundance changes during anoxia.** Proteins were clustered based on log_2_ fold changes (LFCs) across four time points (T1, T2, T4, T24) for two comparisons: anoxic versus dark-oxic conditions (left panel, A*vs*O) and anoxic versus the initial light-oxic control (right panel, A*vs*C). Colors indicate relative abundance from lower (blue) to higher (red). Each condition includes four independent biological replicates. Cluster sizes are indicated next to each group.

To illustrate the overall distribution of abundance changes, log2 fold changes (LFCs) from the 24h anoxic versus light-oxic comparison (T24, A*vs*C) were plotted (Supporting information Fig. S 1). The near-symmetric distribution of fold changes reveals a broad yet balanced proteomic reprogramming under anoxia, rather than a unidirectional stress response.

Protein abundance profiles (Table S1) were grouped by hierarchical clustering for the anoxic versus control comparison (A*vs*C) and for the anoxic versus dark-oxic comparison (A*vs*O) (Fig. 1b). Differential regulation was quantified using a LFC threshold of > 0.25 for increased abundance and <-0.25 for decreased abundance (A*vs*O or A*vs*C). This analysis resolved several distinct temporal patterns of regulation across the four time points (T1, T2, T4, T24). In the A*vs*O analysis, clusters 8, 9 and 10 (124, 14 and 6 proteins, respectively) showed a marked increase in abundance after 24h of anoxia, whereas clusters 3, 4, and 5 contained proteins that were consistently less abundant under anoxia at this time point (Fig. 1b).

Proteins showing a consistent increase in abundance under anoxia were identified using the following criteria: LFC > 0.25 for both A*vs*C and A*vs*O comparisons, and LFC < 0.25 O*vs*C. Using these thresholds, no proteins showed increased abundance after 1h of anoxia. In addition, a progressive increase of proteins number is observed when stress conditions are longer. This delayed response indicates that metabolic reprogramming is progressive and primarily established during prolonged oxygen deprivation (Table 1).

**Table 1.** Number of upregulated and downregulated proteins over time under anoxic conditions. Proteins that were upregulated (LFC > 0.25) or downregulated (LFC < -0.25) in *Thalassiosira pseudonana* under anoxic conditions compared to control conditions (A*vs*C) and oxic conditions over time (A*vs*O) at 1h, 2h, 4h and 24h. The total count represents the cumulative number of regulated proteins across all time points. The percentages represent the proportion of regulated proteins at each time point relative to the total number of upregulated or downregulated proteins within each comparison category.

| *Proteins count* |  |  |  |  |  |
| --- | --- | --- | --- | --- | --- |
| *Conditions* | 1 h | 2 h | 4 h | 24 h | Total |
| *AvsC LFC > 0.25* | 55 (11.4 %) | 114 (23.6 %) | 106 (22.0 %) | 207 (43.0 %) | 482 (100 %) |
| *AvsC LFC < -0.25* | 58 (13.8 %) | 76 (18.0 %) | 125 (29.7 %) | 162 (38.5 %) | 421 (100 %) |
| *AvsO LFC > 0.25* | 47 (14.3 %) | 43 (13.1 %) | 57 (17.3 %) | 182 (55.3 %) | 329 (100 %) |
| *AvsO LFC < -0.25* | 79 (22.2 %) | 54 (15.2 %) | 76 (21.3 %) | 147 (41.3 %) | 356 (100 %) |

Six proteins fulfilled the criteria after 2h, ten after 4h and forty-nine after 24h (two of which correspond to indistinguishable MS sequences and thus appear as only one sequence).

In total, 64 proteins showed specific increased abundance under anoxia, representing < 7% of the detected proteome. Among them, 48 proteins were annotated (3 at T2, 8 at T4 and 37 at T24) (Fig. 2), proportions consistent with the annotation frequency in the complete dataset (Martin, 2015; Wood *et al*., 2019; Plouviez & Dubreucq, 2025).

**Figure 2.**
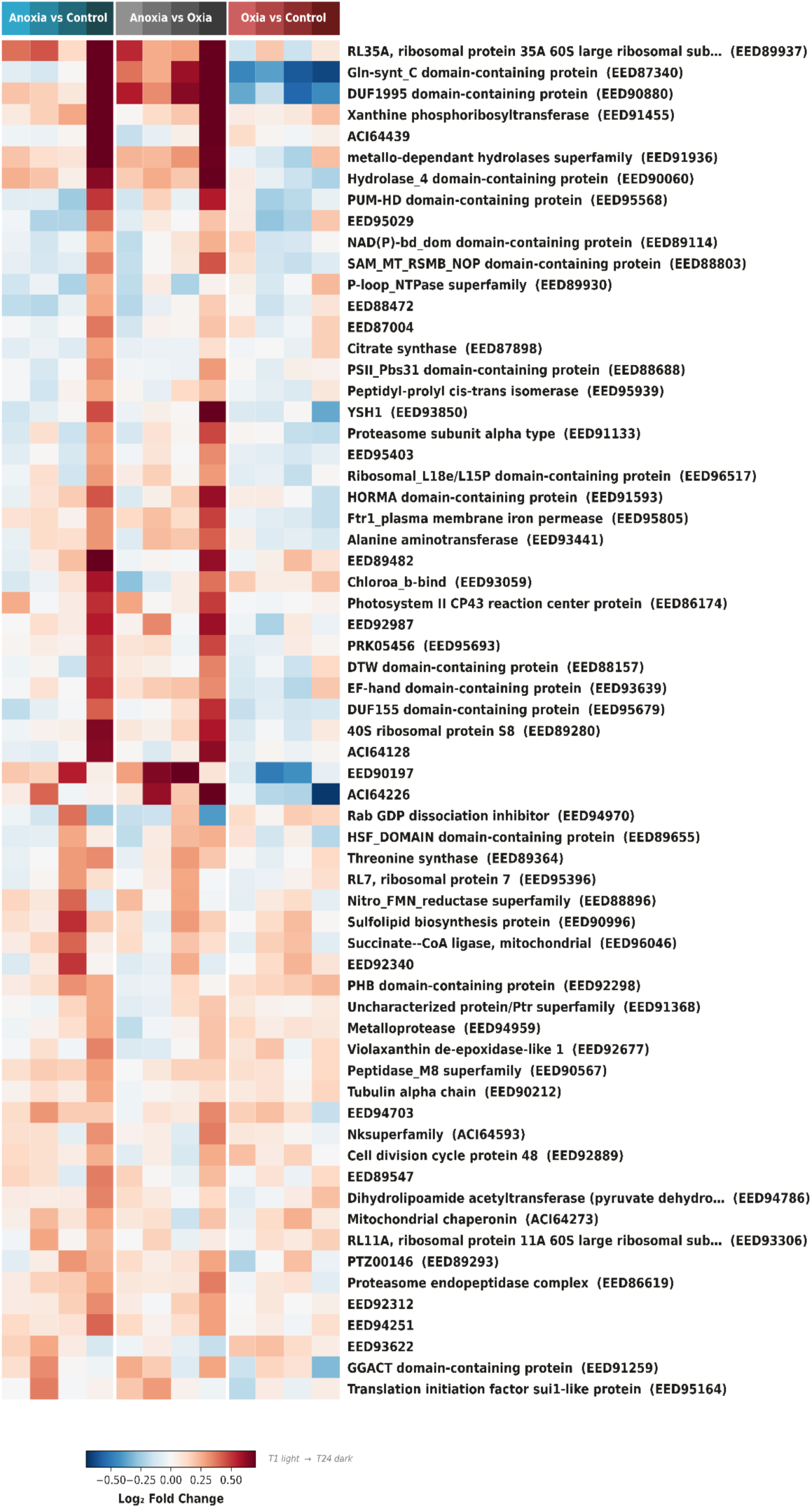
Differentially abundant proteins under anoxic conditions. Heatmap of proteins meeting the following criteria: log2 Fold Change (LFC) > 0.25 for anoxia *vs* control (A*vs*C) and anoxia *vs* oxic (A*vs*O, and LFC < 0.25 for oxic *vs* control (O*vs*C). Protein abundance changes are shown across four time points (T1: 1h; T2: 2h, T4: 4h, T24: 24h). Colors range from blue (lower abundance) to red (higher abundance). Annotated and uncharacterized proteins are included. Proteins listed comprise functions related to central metabolic, amino acid biosynthesis, photosynthesis, protein turnover, and stress-associated processes.

Several proteins increase in abundance both under anoxic and oxic dark conditions (Fig. 1b), indicating that some pathways active during anoxia are not strictly anoxia-specific but are nevertheless likely involved in the overall cellular adjustment. For this reason, the following descriptions of the results focus more broadly on changes in protein abundance based on the anoxia versus control comparison (A*vs*C), with particular emphasis on the 24h anoxic condition, which displays the largest number of proteins with increased abundance (Fig. 1, Fig 2). This approach enables a more comprehensive description of the metabolic pathways engaged during anoxia and provides a basis for interpreting their functional contribution.

Among the 49 proteins whose abundance increased after 24h of anoxia, several are associated with major cellular processes, including amino acid metabolism (glutamine synthase [GLNN], threonine synthase, alanine aminotransferase [ALAT1], the tricarboxylic acid (TCA) cycle (citrate synthase [CSN1], succinate-Coa ligase [SCS1]), the cell cycle (cell division cycle protein 48), photosynthesis (PSII PSBC, PSII Psb31 (Supporting Information Note S1)), and stress-associated pathways (*e.g.* proteasome subunit, proteasome endopeptidase complex) (Note S1) (Fig. 2).

### Alanine accumulation alongside glutamine and aspartate depletion reveals a central metabolic shift linking carbon and nitrogen metabolism under anoxia

Since amino acid metabolism emerged as one of the most represented functional categories in the proteomic dataset (Fig. 1a), intracellular amino acids were quantified under anoxic and oxic dark conditions at 2h and 24h. Principal component analysis (PCA) separated the 24h anoxic condition from all other treatments, indicating a distinct metabolic state associated with prolonged oxygen deprivation (Fig. 3a). In contrast, early anoxic samples partially overlap with oxic dark samples, whereas the light oxic control formed a distinct cluster.

**Figure 3.**
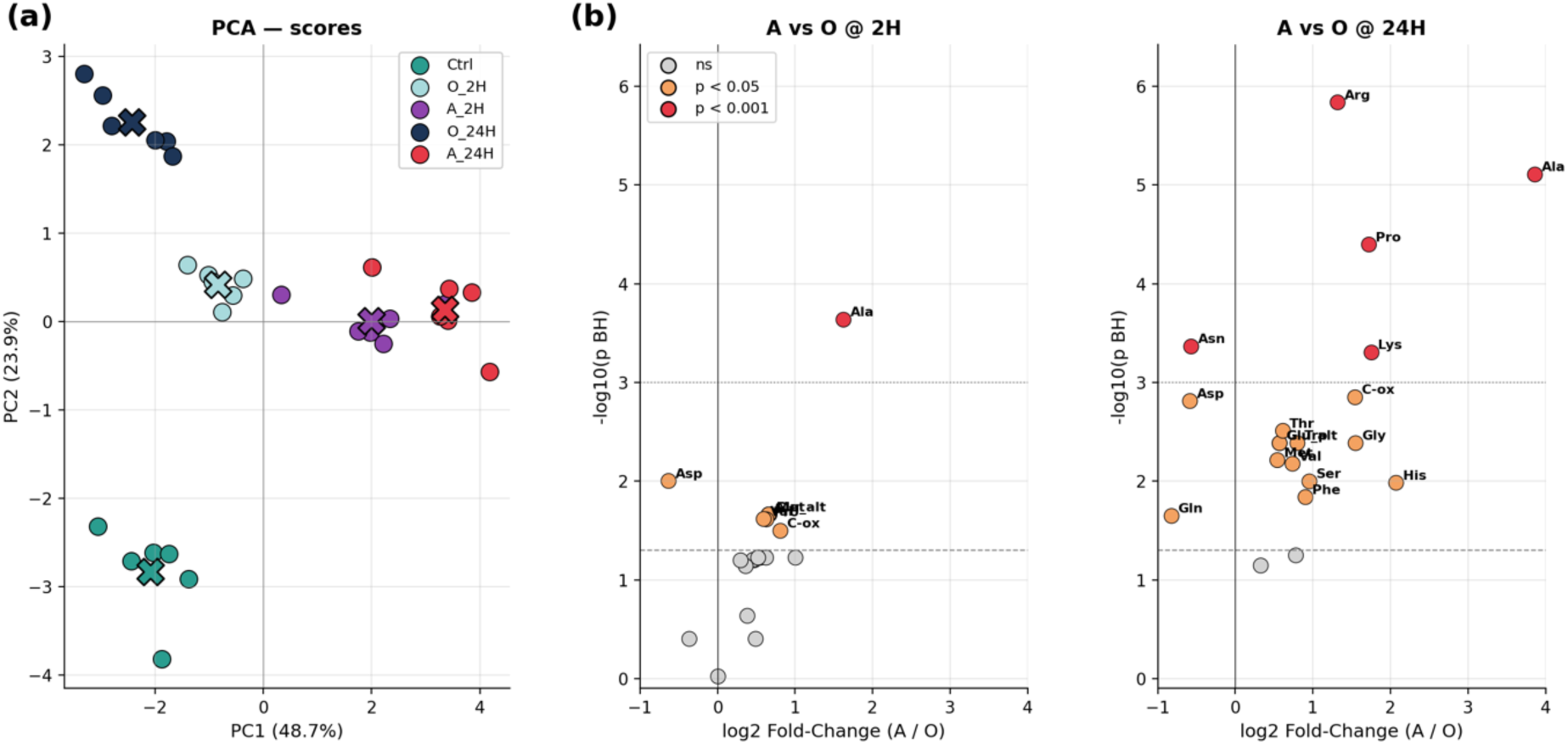
Amino acid composition in *Thalassiosira pseudonana* under anoxic and oxic dark conditions. **(a)** Principal component analysis (PCA) of amino acid profiles under anoxic (A) or oxic (O) dark conditions at 2 h and 24 h, compared to the light-oxic control (Ctrl). each condition includes six biological replicates, represented by colored dots. **(b)** Log2 fold change (LFC) of each amino acid is shown for anoxic versus oxic conditions after 2 h (AvsO @2 h) and 24 h (AvsO @24 h) of treatment. Gray dots indicate non-significant changes, orange dots indicate p<0.05, and red dots indicate p<0.001. Alanine (Ala) shows a significant increase under anoxia, already after 2 h of anoxic exposure, using a on a one-way ANOVA with FDR correction for six biological replicates.

A marked accumulation of methionine was observed under both darkness and anoxia comparing to control condition at both time exposure (O_2H vs Ctrl, A_2H vs Ctrl, O_24H vs Ctrl, A_24H vs Ctrl) (Fig. S2), reaching levels nearly tenfold higher than in the light condition. This response, being common to both treatments, suggests that it is not specific to oxygen deprivation but rather reflects a broader metabolic adjustment associated with the absence of photosynthetic activity. Methionine is a central component of sulfur and one-carbon metabolism, serving as the precursor of S-adenosylmethionine, a universal methyl group donor involved in numerous biosynthetic and regulatory processes (Hanson & Roje, 2001; Hesse *et al*., 2004). Its accumulation may therefore result from a reduced demand for protein synthesis and methylation reactions under energy-limited conditions, leading to an imbalance between synthesis and utilization.

Alanine exhibited the most pronounced increase, particularly after 24h of anoxia (LFC = 4, A *vs* O) (Fig. 3b), with a detectable rise already at 2h (LFC > 1.5). and high confidence in both conditions *(p < 0.001)*. In contrast, glutamine (LFC < -2), aspartate (LFC < -0.9) were consistently depleted under anoxia (A_24H vs Ctrl), while remaining stable showing no significant change under oxic dark conditions (O_24H vs Ctrl) (Supplementary Data S2). Glutamine, aspartate, and asparagine were likewise depleted under anoxia compared with the oxic condition at 24h (A vs O @ 24H) (Fig. 3b). This reciprocal pattern points to a selective redistribution of carbon and nitrogen fluxes rather than a general metabolic slowdown. The concomitant depletion of key nitrogen-rich amino acids further suggests a large-scale reorganization of nitrogen metabolism.

### Carbon metabolism is maintained under anoxia but redirected toward alternative end products

Changes in amino acid and nitrogen metabolism are tightly connected to central carbon pathways. Proteomic data indicate that central carbon metabolism remains active under anoxia, with most glycolytic enzymes detected and only moderate changes in abundance (Supplementary Data table 1, 2), including fructose bisphosphate aldolase (ALDO2), phosphoglycerate kinase (PGK), enolase (ENO1), glyceraldehyde-3-phosphate dehydrogenase (GAPC3), pyruvate kinase (PYK4), triose-phosphate isomerase (TPI3), and phosphoglycerate mutase (PGAM2). This suggests that glycolytic flux is maintained despite the absence of oxygen. Consistently, only limited variations were observed across the pathway, with a modest increase in enolase (ENO1, LFC = 0.33, A*vs*C) and the E2 component (DLAT) of the pyruvate dehydrogenase complex (LFC = 0.35, A*vs*C), and a decrease in phosphoglycerate mutase (PGAM2, LFC = -0.28, A*vs*O) after 24 h, indicative of fine-tuning rather than large-scale reprogramming (Fig. 2, Table S1, Table S3).

In contrast, the tricarboxylic acid (TCA) cycle displayed a more heterogeneous response. Enzymes with increased abundance include citrate synthase (CSN1, LFC = 0.29 at T24, A*vs*C), two isocitrate dehydrogenase (EED93639, LFC = 0.52 at 24h, A*vs*C; EED95879, LFC = 0.36 at 4h, A*vs*C), and two succinate dehydrogenase (SDH1; EED93895, LFC = 0.44 at 24h, A*vs*C; and EED89006, LFC = 0.33 at 24h, AvsC). In contrast, aconitase (acnB; LFC = -0.25 at 24h, A*vs*O) and malate dehydrogenase (MDH1; EED95757) (LFC = -0.31 at 24h, A*vs*O), showed decreased abundance, a pattern consistent with a reduced reliance on fully oxidative metabolism.

Under these conditions, the maintenance of glycolysis is expected to generate excess reducing equivalents that cannot be reoxidized via mitochondrial respiration. Several enzymes potentially involved in alternative redox-balancing pathways showed increased abundance, including a succinyl-CoA ligase (SCS1, LFC = 0.41 at 4h, A vs C), a NAD-dependent malic enzyme (MAO1, LFC = 0.55 at 24h, A *vs* O), a predicted alcohol dehydrogenase (EED94859, LFC = 1.43 at 24h, A *vs* C), and a predicted butyryl-CoA dehydrogenase (EED89520; LFC = 0.43 at 24h, A *vs* C). Besides, Pyruvate formate lyase (PFL) showed time-dependent changes, increasing at 1h (LFC = 0.33, A *vs* O) and decreasing at 24h (LFC = -0.68, A *vs* C). Alanine production by ALAT provides an additional route for pyruvate utilization that avoids further NADH generation. Finally, lactate dehydrogenase (LDH) and acyl-CoA ligase (ACS1), two enzymes typically involved in fermentative metabolism in *Chlamydomonas reinhardtii* (Atteia *et al*., 2013) showed no detectable changes in abundance in *T. pseudonana* under any of the conditions tested.

The pentose phosphate pathway also showed selective modulation, with upregulation of two enzymes from the non-oxidative branch, namely ribose-5-phosphate isomerase (EED94748) and a transaldolase (TAL) (EED87192; EED87575), suggesting a role in carbon redistribution and biosynthetic processes.

Importantly, multiple metabolic connections link nitrogen and carbon metabolism. The urea cycle generates fumarate, which can feed into the TCA cycle, while transamination reactions connect glutamate to central carbon intermediates. These interactions support a flexible metabolic network that maintains energy production and redox balance under anoxia (Fig. 5).

### Proteomic data revealed extensive modulation of nitrogen-related pathways under anoxia

Key components of the DNRA pathway were detected in the dataset (Fig. 2). These include NADPH nitrate reductase (NIR1), nitrite reductase (NIR2), and a predicted nitrite reductase (EED90961), the latter showing a marked increase in abundance after 24h of anoxia (LFC = 2.33, A*vs*C; LFC = 0.77, A*vs*O). A nitrate transporter (NRT1) also displayed increased abundance at 24h (LFC = 0.36, A*vs*O). However, most DNRA-associated activities were present under both oxic and anoxic conditions, indicating that these changes are not specific to anoxia.

In parallel, enzymes of the urea cycle were also detected. Especially, enzymes from the linear pathway (Allen *et al*., 2011; Winter *et al*., 2015) were either present and constant or upregulated. While carbamoyl phosphate synthase (CPS1, EED92873) and urease did not display notable changes in abundance, two enzymes showed increased abundance either in anoxia or in darkness at 24h: argininosuccinate synthase (ASS, ACI64235) increased specifically under anoxia (LFC = 1.56, A*vs*C; 0.64, A*vs*O), while ornithine carbamoyltransferase (OTC, EED96227) increased in both dark conditions (LFC = 0.39, AvsC). Notably, this part of the cycle forms arginine and fumarate from aspartate. In contrast, N-acetyl-gamma-glutamyl-phosphate reductase (AGPR), involved in the so-call urea cyclic pathway (Winter *et al*., 2015) is downregulated (EED94749, LFC = -0.37, A*vs*C; -0.5 A*vs*O).

In addition to ALAT1, another prominent feature of this response is the strong increase of glutamate ammonia ligase (GSIII), at all time points under anoxia (LFC = 1.44, A*vs*O; 0.80, A*vs*C at 24h), suggesting active ammonium assimilation. At the same time, the depletion of glutamine implies rapid turnover and redistribution of nitrogen.

Together, these data indicate that nitrogen metabolism is not merely maintained under anoxia but reorganized into a central hub that supports amino acid interconversion, redox balance, and metabolic flexibility.

Importantly, these metabolite changes are consistent with the proteomic data. Alanine aminotransferase (ALAT1), which catalyzes the conversion of pyruvate and glutamine into alanine and alpha-ketoglutarate, showed increased abundance under anoxia (LFC = 0.31, A*vs*C; 0.42, A*vs*O at 24h). This supports a model in which pyruvate derived from glycolysis is redirected toward alanine production, while glutamate serves as an amino donor, linking carbon metabolism to nitrogen assimilation.

This apparent imbalance is consistent with increased demand for glutamate, a central metabolic node linking nitrogen and carbon metabolism. Supporting this view, xanthine phosphoribosyltransferase, a rate-limiting enzyme in purine metabolism (Roy *et al*., 2015; Glockzin *et al*., 2023), was strongly upregulated (LFC = 1.22, A*vs*C; 1.01, A*vs*O). Its activity produces xanthosine monophosphate (XMP) from xanthine and 5-phospho-d-ribose 1-pyrophosphate, providing a potential route for glutamate production.

### The anoxic response combines conserved functions with lineage-specific innovations

To investigate the evolutionary basis of this response, we analyzed the phylogenetic distribution of the 64 anoxia-responsive proteins. Most proteins are broadly conserved across Stramenopiles, with subsets found restricted to Gyrista, Ochrophyta, Diatomista, and Bacillariophyta (Fig. 4, Fig. S1). Proteins shared among all Stramenopiles constitute the largest group which include several proteins (*e.g*. Succinyl-CoA synthetase beta chain [SCS1]) contributing primarily to the early response phase (T2 and T4) and representing over a quarter of the responsive proteins. These proteins include alanine aminotransferase (ALAT), involved in alanine production, conserved in all Stramenopiles.

**Figure 4.**
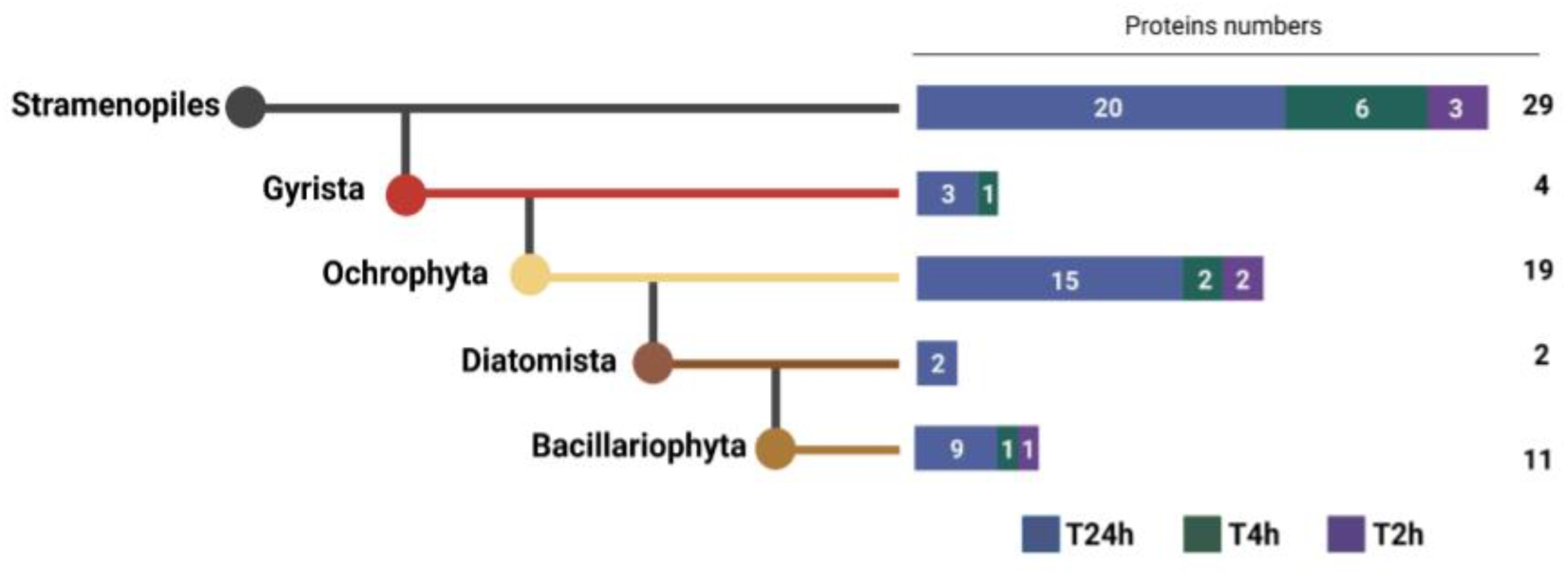
Phylogenetic distribution of proteins associated with the anoxia response in *Thalassiosira pseudonana*. Proteins identified as responsive to anoxia were assigned to phylogenetic groups based on their evolutionary distribution. The largest category comprises proteins found across all Stramenopiles (dark grey), with additional subsets restricted to Ochrophyta (red), Bacillariophyta (beige), Gyrista (dark brown), and Diatomista (light brown). For each phylogenetic group, bar plots indicate the number of proteins whose abundance changes at 2h (T2, purple), 4h (T4, green), and (T2 in grey and T4 in light orange) and 24h (T24, blue).

In contrast, several other proteins associated with nitrogen and amino acid metabolism display lineage-specific distributions, notably within Gyrista, like threonine synthase (TS, restricted to Gyrista). Other identifiable pathways involved in anoxic stress response had an origin that could be traced back to the divergence of Ochrophyta and plastid acquisition. This is the case of xanthine phosphoribosyltransferase involved in nucleotide metabolism (sup. table 2), which is one of the three highest upregulated proteins at T24 in A*vs*C (LFC=1.22 A*vs*C).

Finally, some proteins involved in anoxic stress responses have been gained more recently in the evolution as glutamine synthase (GSIII, restricted to Diatomista) and gamma-glutamyl cyclotransferase (GGCT, restricted to Bacillariophyta). This combination of conserved and lineage-specific components indicates that the anoxic response in *T. pseudonana* results from the integration of an ancient metabolic framework with more recent innovations, likely enhancing metabolic flexibility in low-oxygen environments.

## Discussion

### An integrated metabolic response to anoxia centered on amino acid and nitrogen fluxes

Diatoms are frequently encountered in low-oxygen environments (Muylaert *et al*., 2006; Broman *et al*., 2017), yet the metabolic basis of their survival under anoxia has remained poorly understood. Here, by integrating proteomic, metabolomic, and phylogenomic analyses, we show that the centric diatom *Thalassiosira pseudonana* responds to prolonged anoxia through a coordinated reorganization of central metabolism, in which amino acid and nitrogen fluxes play a central role.

A key finding of this study is the strong and specific accumulation of alanine under anoxia, accompanied by the depletion of glutamine, aspartate, and asparagine, indicating a major reorientation of both carbon and nitrogen toward alanine synthesis. The concomitant increase in alanine aminotransferase (ALAT1) abundance, together with the marked rise in alanine levels in metabolomic assays, strongly supports the existence of an alanine-centered fermentative metabolic route in *T. pseudonana* (Fig. 2, 3b, and 5). Alanine content increased particularly after 24h of anoxia (LFC = 3, A*vs*C), making it one of the most responsive amino acids under these conditions.

This accumulation likely results from transamination reactions between pyruvate and glutamate, providing an efficient way to consume pyruvate without generating additional NADH generation, and thereby contributing to redox balance when mitochondrial respiration is impaired. In this context, alanine production from glutamate and pyruvate may act both as a nitrogen sink and as a mechanism for metabolic adjustment under oxygen deprivation (Fig. 5).

**Figure 5.**
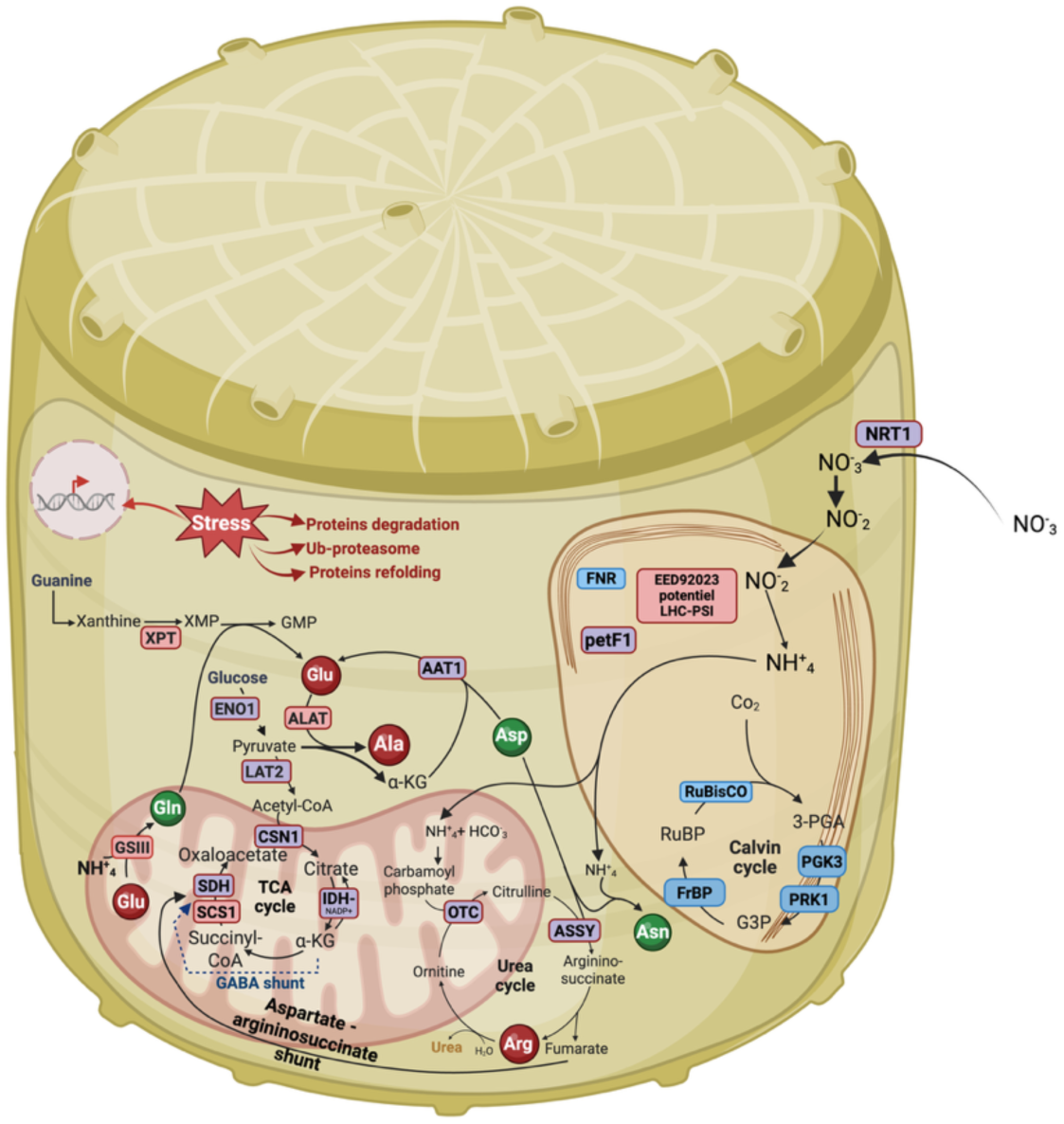
Overview of metabolic adjustments in *Thalassiosira pseudonana* under dark anoxia. Schematic representation of the major metabolic pathways affected during dark anoxia in *T. pseudonana*, integrating proteomic and metabolomic data. Proteins highlighted in light blue show increased abundance specifically in darkness (under both oxic and anoxic conditions), whereas proteins in red show increased abundance specifically in anoxia. Amino acids in red display increased intracellular levels under anoxia, while those in green show decreased levels. The diagram summarizes changes across nitrogen metabolism (including DNRA and the urea cycle), glycolysis, the TCA cycle, putative fermentative routes, and stress-associated processes, illustrating the coordinated metabolic adjustments supporting anaerobic acclimation.

Although alanine accumulation has been well documented in plant roots upon hypoxia stress (Miyashita *et al*., 2007; Rocha *et al*., 2010; Diab & Limami, 2016) and also reported in some anaerobic non-photosynthetic eukaryotes such as *Entamoeba histolytica* or *Trichomonas vaginalis* (Catalanotti *et al*., 2013), it is not generally considered a dominant feature of anaerobic metabolism in photosynthetic cells. In the green alga *Chlamydomonas reinhardtii*, alanine accumulation under anoxia remains comparatively limited (LFC = 1) (17) and appears to become especially pronounced in mutant strains lacking functional pyruvate formate lyase (PFL) (LFC > 3; (18)), suggesting that alanine fermentation is not a major route in that species. The pronounced alanine response observed here therefore reveals a distinct metabolic strategy in the diatom *T. pseudonana*, in which alanine fermentation appears to be a central component of the anoxic response.

### Central Role of Nitrogen Metabolism

Both our proteomic and metabolomic data further support nitrogen metabolism as a cornerstone of the anoxic response (Kamp *et al*., 2011, 2015, 2016). The complete urea cycle was retained under anoxia, with increased abundance of enzymes such as ornithine carbamoyltransferase (OTC) and argininosuccinate synthase (ASS), likely ensuring nitrogen conservation and redistribution (Hockin *et al*., 2012). The DNRA pathway, known to support intracellular nitrate reduction during anoxia (Kamp *et al*., 2011, 2015; Stenow *et al*., 2024), appears to remain active even after extended exposure, as indicated by the upregulation of NIR2 at 24h (LFC = 2.3). The strong upregulation of glutamine synthetase (GSIII) combined with the depletion of glutamine, suggests a high metabolic demand for glutamate, and ammonium assimilation under anoxia, mirroring observations in the green alga *Selenastrum minutum* (Vanlerberghe & Turpin, 1990).

Because alanine production by ALAT requires glutamate consumption, maintaining glutamate availability under anoxia is likely important. In this context, glutamate appears to occupy a central position at the intersection of several pathways, including GSIII activity, alanine synthesis, and possibly other metabolic rearrangements, despite its relatively stable abundance over time in targeted metabolomics (Fig. 4 and 5). By contrast, glutamine levels clearly decreased, suggesting active conversion into other metabolites. One possible contributing route is purine metabolism, as xanthine phosphoribosyltransferase, which converts xanthine to XMP, was specifically more abundant under anoxia (Fig. 2). Because XMP can subsequently be used in reactions involving glutamine and leading to GMP production with glutamate release via GMP synthase activity (Fig. 5 and Table S 3), this pathway could contribute to glutamate regeneration under anoxia. Such a mechanism has been described in hypoxic human cells (Yoo *et al*., 2020), although its role in *Thalassiosira* remains hypothetical. Interestingly, guanine-rich storage structures have been reported in several algae and protists (Mojzeš *et al*., 2020; Pilátová *et al*., 2022; Goodenough *et al*., 2025), raising the possibility that intracellular purine reserves could support this metabolism, although this remains entirely speculative in the present context. Overall, these observations suggest that purine metabolism may participate in metabolic adjustment to anoxia, but this will require direct experimental validation.

### Metabolic Flexibility via Fermentative Pathways

Seven percent of the detected proteins were specifically upregulated in anoxic conditions (excluding responses to darkness alone), suggesting a tightly regulated and targeted response to oxygen limitation. Among identified proteins, several enzymes predicted to function in fermentative metabolism (Atteia *et al*., 2013) were confirmed, including PFL, LDH, MAO1, NIR1/NIR2 and ACS1 (Table S 1). Proteins such as MAO1, succinyl-CoA ligase (SCS1), pyruvate formate lyase (PFL), and a predicted alcohol dehydrogenase displayed time-dependent abundance changes, supporting the coexistence of multiple, non-exclusive routes for pyruvate utilization and redox balancing. Interestingly, enzymes potentially associated with butyrate metabolism, such as succinyl-CoA ligase and lactate dehydrogenase, were upregulated (Fig. 2 and Table S3). Since the proteins showing increased abundance specifically under anoxia represent only a small subset of the cellular response, this may indicate, beyond alanine fermentation, a broader fermentative strategy than previously documented in diatoms.

### Sustained glycolysis and partial TCA cycle remodeling

During anoxia, enzymes of the pentose phosphate pathway (PPP), notably TAL and RPI, were more abundant at all time points (Fig. 2 and Table S3). By contrast, glycolytic enzymes did not show any strong variations and remained abundant throughout the stress, with ENO1 among the most abundant proteins detected. This pattern suggests that glycolysis is maintained under anoxia, while the PPP may act as a parallel branch, possibly supporting both carbon rerouting and nucleotide-related metabolism, consistent with the high abundance of XPT. In parallel, pyruvate may still be directed toward acetyl-CoA production by LAT2 and subsequently feed the TCA cycle.

TCA cycle remodelling is a common feature of anaerobic metabolism because some of its steps are tightly linked to respiration (Vanlerberghe *et al*., 1989; Martínez-Reyes & Chandel, 2020). In our data, SCS1 and IDH1 were more abundant at 4h but returned to control levels by 24h, suggesting that a more conventional TCA cycle may operate at early stages of oxygen depletion, whereas prolonged anoxia favors a more partial functioning of the cycle (Table S1 and Table S 3). Under these conditions, metabolic shunts may help maintain TCA intermediates while bypassing reactions normally associated with oxidative phosphorylation (Fig. 5).

A first shunt candidate is the urea cycle, already recognized as a central metabolic hub in diatoms (Allen *et al*., 2011; Prihoda *et al*., 2012). Urea cycle enzymes remain highly abundant during anoxia, especially at the late stage, and this pathway can supply fumarate through argininosuccinate lyase, as previously proposed by Allen et al. (Allen *et al*., 2011). Its potential contribution is also noteworthy because of its connection with nitrogen metabolism via DNRA, a process central to anaerobic metabolism in diatoms (Kamp *et al*., 2011, 2015, 2016). Consistent with this idea, CPS abundance remained stable across conditions, supporting the possibility that NH4+ derived from DNRA could continue to feed carbamoyl phosphate production for the urea cycle (Fig. 5). Another possible alternative is the GABA shunt, previously described in *P. tricornutum* (Matthijs *et al*., 2017), which may also help maintain TCA intermediates under prolonged anoxia. Altogether, these results support a metabolic shift under anoxia in which sustained glycolysis and PPP activity are associated with a transition from a more canonical TCA cycle during early stage of anoxia to a more partial configuration under prolonged anoxia, potentially supported by the urea cycle and GABA shunts.

### Evolutionary perspective on diatom anaerobic metabolism

Chimeric genomes of diatoms (Ochrophyta) derived from secondary plastid endosymbiosis and massive lateral gene transfers from various sources (Allen *et al*., 2006; Archibald, 2009, 2015; Dorrell *et al*., 2017, 2023). The phylogenetic distribution of the 64 specifically upregulated proteins reveals that while early anoxic responses are driven by conserved Stramenopile genes, some specific functions are restricted to Diatomista and Bacillariophyta (Fig. 4). This suggests more recent gene acquisition putatively linked to LGT, as it has been shown for various adaptation in Stramenopile organisms (Allen *et al*., 2011; Eme *et al*., 2017; Dorrell *et al*., 2023). However, this enzyme is restricted to Diatomista (Table S2, Fig. S3), suggesting specific response of this group of Algae. Moreover, the presence of enzymes like gamma-glutamyl cyclotransferase (GGCT) specific to Bacillariophyta implies lineage-specific mechanisms for maintaining nitrogen homeostasis, possibly acquired through lateral gene transfer (LGT), as already described in diatoms to adapt to other environments (Dorrell *et al*., 2023). This activity is linked to the glutathione cycle, a crucial metabolism in hypooxia/anoxia in animals (Taniguchi *et al*., 2022; Tabata *et al*., 2024) and in other organisms (Mojzeš, 2025), such as plants under stress conditions (Ristova & Kopriva, 2022). Interestingly it seems to be linked to amino acid supplies in various cases.

LGT have provided diatoms with metabolic toolkits enabling them to thrive in fluctuating environments (Allen *et al*., 2011; Dorrell *et al*., 2017, 2023). The phylogenomics analysis reveals that the anoxic response combines conserved eukaryotic functions with lineage-specific innovations. Core metabolic pathways involved in central carbon metabolism (*e.g.* CSN1 or isocitrate dehydrogenase from the TCA cycle) early responses are widely conserved across Stramenopiles, indicating an ancient metabolic framework for coping with oxygen limitation. As well, ALAT enzyme is found in all Sramenopiles, while being central to generate from glutamine, alanine and alpha-KG and coupling carbon metabolism to nitrogen assimilation, as discussed above, and is likely only a readaptation of ancestral metabolism to cope with anoxic conditions.

Moreover, xanthine phosphoribosyltransferase can be traced back to Ochrophyta, therefore could be linked to acquisition of plastid. In contrast, several key components of nitrogen and amino acid metabolism, including glutamine synthetase (GSIII), already described in Allen et al. 2006 (Allen *et al*., 2006), and gamma-glutamyl cyclotransferase (GGCT) have been acquired during or after the divergence of the diatomista. The recruitment of GSIII could be linked to the well-known DNRA process (Fig. 5), and highlights how LGTs have enhanced a likely, already existing anaerobic gene repertoire. The integration of these ancient conserved and more recently acquired components results in a chimeric metabolic network that enables efficient adaptation to fluctuating oxygen conditions (Fig. 5). This evolutionary layering may help explain the ecological success of diatoms in environments characterized by recurrent or prolonged hypoxia.

## Conclusion - A metabolic framework for anoxia tolerance in diatoms

Our results support a model in which anoxia tolerance in *T. pseudonana* does not rely on a single metabolic pathway, but rather on the coordinated integration of multiple processes. Central to this response is the reorganization of amino acid and nitrogen metabolism, which supports carbon flux redistribution, redox balance, and metabolic flexibility.

In this framework, alanine production emerges as a key metabolic output, while glutamate functions as a central hub linking nitrogen and carbon metabolism. The coupling of these processes with partial TCA activity and metabolic shunts allows the maintenance of energy production under oxygen limitation.

These adaptive strategies reflect both ancient, conserved responses and more recent, lineage-specific innovations. This integrated metabolic strategy differs from those described in other photosynthetic eukaryotes and highlights the unique adaptations of diatoms to low-oxygen environments. More broadly, these findings provide a conceptual framework for understanding how phytoplankton can persist expanding oxygen minimum zones in modern oceans.

## Acknowledgement

This work was supported by the Belgian Fonds de la Recherche Scientifique (FNRS) (grants nos PDR T.0032, and J.0025.24 to PC). PC is a Research Director of FNRS. P.C. is Senior Research Associate from Fonds de la Recherche Scientifique – FNRS. U.C. acknowledges CNRS to provide him a “delegation” at the University of Liège (ULiège), as well as ANR project CONVERGE, and CDP-PIE

## Author Contributions

G.G., M.C., U.C. and P.C. conceived the research. G.G., M.C., and N.B performed protein extraction for proteomic analysis and metabolomic analysis. H.D. and P.M. performed mass-spectrometry. G.G., M.C, U.C., P.C. generated all figures, and M.C. performed the heatmap (clustering analysis). G.G., M.C., U.C. and P.C. analyzed proteomic data. M.C. and F.B. performed the metabolomic approach, and M.C. and P.C. analyzed it. U.C. and M.C. performed the phylogenetic and localization prediction analysis. M.C., G.G., U.C. and P.C wrote the manuscript. All the authors have reviewed the manuscript and approved the final version.

## Data availability

The mass spectrometry proteomics data have been deposited to the ProteomeXchange Consortium via the PRIDE (Perez-Riverol *et al*., 2025) partner repository with the dataset identifier PXD079012 and 10.6019/PXD079012.

The mass spectrometry supporting amino acids metabolomics are available at Zenodo with DOI: 10.5281/zenodo.19819410

## Supporting Information (brief legends)

**Supporting Information Note S1**. Results and discussion of protein identified by proteomic link to photosynthesis, as well as other metabolism footprint putatively involved.

**Supporting Information Fig. S1**. Distribution of log_2_ fold change (LFC) for the 928 identified proteins

**Supporting Information Fig. S2** Log_2_ fold change (LFC) of amino acids for each condition (Oxic vs Ctrl; Anoxic vs Ctrl; after 2h or 24h of treatment).

**Supporting Information Fig. S3** All phylogenetic trees obtained for the 64 upregulated proteins identified by proteomic are available in Supplementary data 1.

**Supporting Information Table S1.** Table of all proteins detected by proteomic with their annotation and their relative abundance based on the various conditions.

**Supporting Information Table S2.** Table with 64 upregulated proteins identified by proteomic, with their phylogenetic annotation (*i.e.* lineage distribution) as well as their predicted localization.

**Supporting Information Table S3.** Table focused on all proteins discussed in the article, with annotation and their relative abundance based on the various conditions.

