## Supplemental figure 1, 2 for "Proteomic Response of *Thalassiosira pseudonana* to Anoxia Reveals Alanine Fermentation Pathway and Reprogramming of Nitrogen Metabolism"

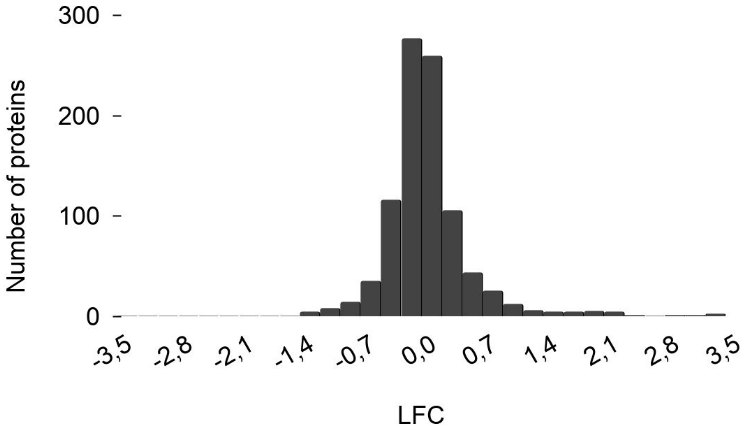


**Supplemental Fig. S 1** : Distribution of $\log_2$ fold change (LFC) for the 928 identified proteins after 24 h of anoxia relative to the light-oxic control (A*vs*C 24 h). The histogram shows a symmetric distribution centered near zero, with proteins displaying both increased and decreased abundance.

**Figure 1. Overview of the proteomic dataset for *Thalassiosira pseudonana* under anoxia and clustering analysis of protein abundance changes during anoxia.**

**Figure 2. Proteins showing increased abundance under anoxic conditions.**

**Figure 3: Amino acid composition in *Thalassiosira pseudonana* under anoxic and oxic dark conditions.**

**Figure 4. Phylogenetic distribution of proteins associated with the anoxia response in *Thalassiosira pseudonana*.**

**Figure 5: Overview of metabolic adjustments in *Thalassiosira pseudonana* under dark anoxia.**

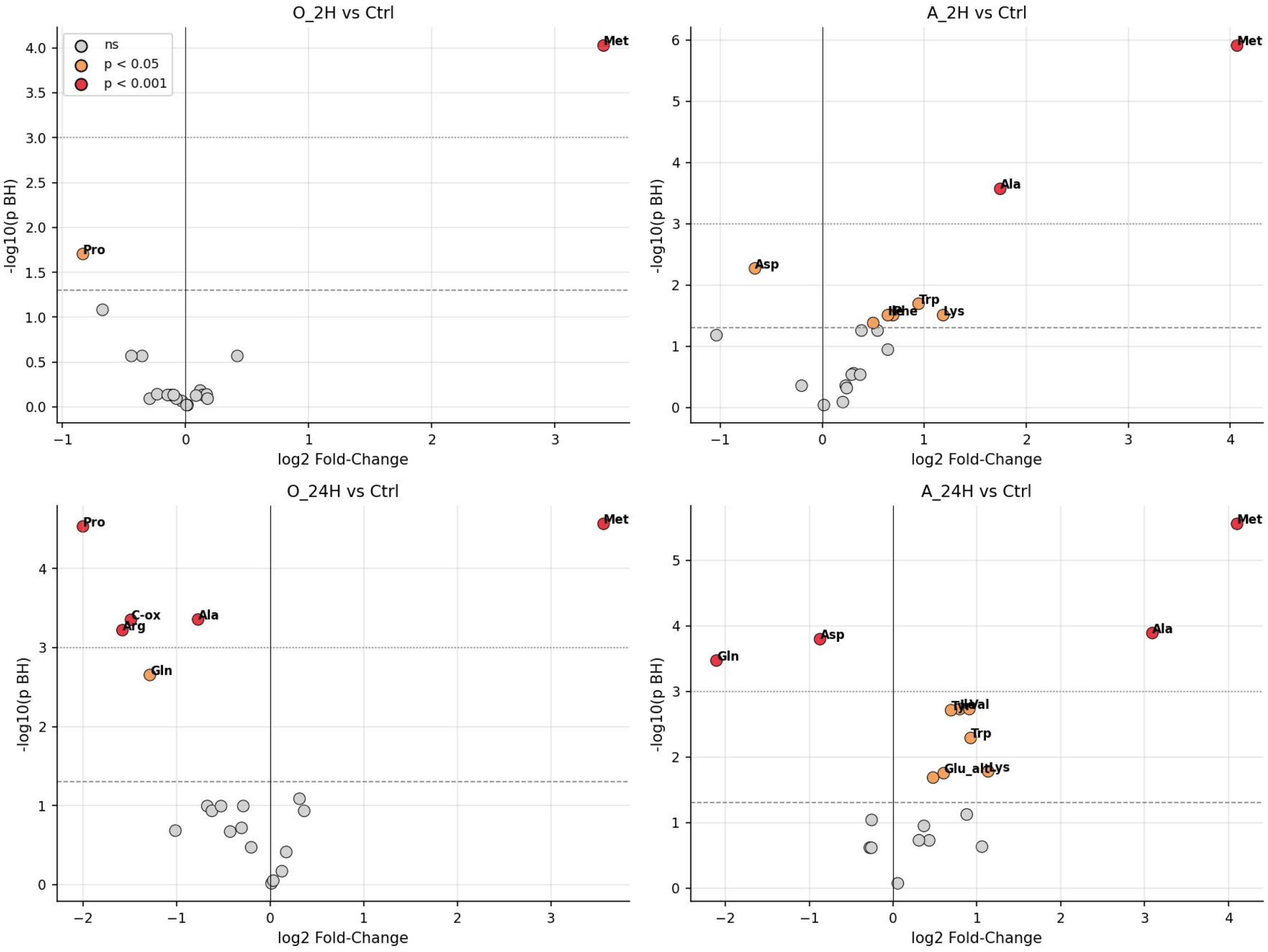


**Supplementary Data S 2** : Log2 fold change (LFC) of each amino acid is shown for oxic versus control conditions after 2 h (O_2H vs Ctrl) and 24 h (O_24H vs Ctrl), and for anoxic versus control conditions after 2 h (A_2H vs Ctrl) and 24 h (A_24H vs Ctrl), based on a one-way ANOVA with FDR correction for multiple testing. Gray dots indicate non-significant changes, orange dots indicate $p<0.05$, and red dots indicate $p<0.001$ (FDR-adjusted p-values)
